# Hierarchical Coarse-to-Fine cGAN for Subtype-Specific Freezing of Gait Signal Generation

**DOI:** 10.1101/2025.10.04.680444

**Authors:** Xinyue Yu, Helena Cockx, Ying Wang, Richard van Wezel, Kaylena Ehgoetz Martens, Arash Arami

## Abstract

Freezing of gait (FOG), a debilitating symptom of Parkinson’s disease, manifests in subtypes as shuffling, trembling, or akinesias, with occurrence and frequency varying across patients. While deep learning (DL) models show promise in FOG detection, their robustness and generalization across subtypes are limited by data scarcity and imbalances between FOG/non-FOG classes and among subtypes. To address this, we propose a subtype-aware FOG augmentation technique enabling training of DL models to perform consistently across subtypes. Specifically, we introduce Hierarchical Coarse-to-Fine conditional Generative Adversary Network (Hi-CF cGAN), a two-stage model that generates subtype-conditioned FOG-like ankle accelerations that are realistic and diverse, as verified through visualization, UMAPs, and Maximum Mean Discrepancy comparison against real signals. We evaluate its effectiveness by training CNNs for FOG detection with both general (subtype-stratified) and personalized (subtype-variant, based on patient-specific subtype composition) augmentation via Hi-CF cGAN, benchmarking against classical augmentations and baseline (no augmentation). Compared to baseline, general augmentation with Hi-CF cGAN effectively improves average detection rates of FOG, trembling FOG, and especially the previously overlooked minor subtypes, shuffling FOG (from 66.8% to 81.6%) and akinesia FOG (from 58.7% to 77.9%). These improvements exceed those of classical augmentations, demonstrating superior realism, richness, and adaptability of Hi-CF cGAN-generated data in addressing FOG/non-FOG and subtype imbalances. Personalized augmentation further enhances accuracy on targeted subtype(s) compared to general augmentation, highlighting its potential for tailored model optimization.

## I. Introduction

**F**REEZING of Gait (FOG) is a brief, episodic loss or freduction of forward movement despite intention to walk [1]. It is a debilitating yet common symptom of Parkinson Disease (PD) that contributes to falls and significantly impacts patients’ quality of life [2]. As a heterogeneous phenomenon, FOG may occur in different medication states and environmental contexts [3]. FOG also varies in manifestations, thereby prompting the definition of gait [4], phenomenological [5], or manifestation-specific [6] subtypes: shuffling-forward (shuffling), trembling in-place (trembling), and akinesia, i.e., complete limb immobility. As subtypes can co-occur within individuals [7], subtype importance may vary across patients based on their subtype compositions and subjective experience: some may prioritize their most frequent subtype(s), while others prioritize the personally most disabling one(s).

To better understand and manage this disruptive and variable phenomenon, many FOG detection algorithms have been developed using wearable sensors, using predominantly kinematic data from inertial measurement units and accelerometers [8]. These algorithms allow FOG progression monitoring, automatic FOG annotation [9], and on-demand activation of FOG intervention or assistive devices [10]. Deep Learning (DL) algorithms, such as feedforward neural network [11], convolutional neural network (CNN) [12], recurrent neural network [13], and transformers [14], demonstrate advantageous detection performances. Handling movement variability via complex pattern capture, they outperform traditional methods like empirical thresholding of the Freezing Index (FI), i.e., power ratio of freeze band (3–8 Hz) to locomotor band (0.5–3 Hz) [15] and classical machine learning models relying on hand-crafted features [16]. However, training a reliable and generalizable DL algorithm requires an extensive [16], diverse [17], and balanced [18] dataset, which is difficult to fulfill in FOG data collections. Specifically, the time-consuming and expertise-required test setup and data post-processing limit dataset size. Considering FOG heterogeneity, limited test protocols and patient sample size in a single collection [19] also restrict dataset diversity. Additionally, difficulty in provoking FOG in a lab or clinical setting [1] results in imbalanced, non-FOG dominated dataset composition [16].

Given collection challenges, typical post-processing for FOG kinematics datasets to cope with data imbalance includes under-sampling of non-FOG [11] [20], segmenting FOG data with higher overlaps [21] [22], assigning higher training weight to the FOG class [23] [24], Synthetic Minority Over-Sampling Technique (SMOTE) (i.e. generating synthetic samples by interpolating between existing, similar ones) or its variations [18] [25], and classical data augmentations (i.e., rotation, jittering, scaling, magnitude warping, permutation, and time warping) [12] [26] [27] [28]. Although undersampling, differential segmentation, and re-weighting help in balancing class size or importance, they do not improve data diversity since they introduce no new information. SMOTE is mainly applied to feature synthesis, as it ignores temporal order within raw signals and may yield unrealistic sequences, limiting its suitability for training DL models that learn from raw data. Classical augmentations improve dataset size, balance, and variability by modifying raw signals [27], but diversity gains are limited by minimal perturbations of real samples [29]. Originally developed for image domain, classical augmentations may also generate unrealistic time-series by distorting original temporal dynamics or magnitude fluctuations. For instance, permutation, i.e., shuffling signal segments, can break the temporal coherence of gait. Magnitude warping, i.e., scaling different signal segments independently, may result in biomechanically implausible amplitude change. Moreover, the optimal augmentation(s) may vary for application goal and sensor setup [27], requiring additional adjustment efforts. Furthermore, as empirically handcrafted heuristics, available augmentations may not align with the characteristics of all signal types. In contrast, directly modeling the data distribution, DL-based data augmentations such as Generative Adversarial Networks (GANs) is more flexible and adaptive, allowing the generation of FOG-tailored signals. Consisting of a generator that synthesize data and discriminator(s) that guide the signal generation via authenticity check, GANs also generates more diverse and realistic FOG signal through its feedback-driven inspector mechanism [29], enabling coverage of a broader motion patterns and FOG characteristics and thereby support the training of more robust prediction models.

Despite its advantages, GAN-based augmentation for FOG kinematics has been explored in only a few studies. Peppes et al. proposed FoGGAN, a GAN generating synthetic FOG signals from 3D accelerometers on shank, thigh, and lumbar region [30]. They demonstrate the similarity of generated data to real data by matching acceleration distributions and across-acceleration correlation matrices. Hadley et al. developed a conditional GAN (cGAN), a GAN variant guided by additional condition, to generate 3D accelerometers signals from lumbar and ankles during FOG or normal PD patient walking [31]. They validated the realism of the synthetic data via its general closeness to real data in visualization (raw signals and signal distribution), FI, and power spectrum, although synthetic data showed slightly lower FI and freeze-band power. While demonstrating the feasibility of realistic FOG generation via GAN, these studies do not assess augmentation impact on DL FOG detection models. Classical augmentation(s) have improved DL classifiers in FOG detection [26] and other PD-related problems (motor state identification between bradykinesia and dyskinesia [28], PD-healthy gait differentiation [27]), motivating investigation into whether DL-based augmentation offers similar or superior benefits despite higher computational cost and complexity. Additionally, FOG heterogeneity, best represented by the composition of manifestation-specific subtypes (hereafter subtypes) with respect to kinematics signals [6], has been neglected by all previous FOG augmentation studies. Subtype-specific FOG generation offers advantages by addressing subtype imbalance, which improves model generalizability across subtypes to account for patient-varying subtype importance. Moreover, without conditioning, the major subtype could dominate synthetic FOG data, deteriorating subtype imbalance. For instance, Hadley et al. observed ‘inactive FOG’ (i.e. flat records like akinesia) dominating their generated FOG data until ‘active FOG’ (gait-pattern signals like shuffling) was explicitly added to the training dataset [31]. Furthermore, subtype-conditioned generation enables customization of detection models to patient’s specific subtype composition, adapting better to FOG heterogeneity.

To tackle FOG heterogeneity which has been overlooked previously, we propose Hierarchical Coarse-to-Fine cGAN (Hi-CF cGAN), an innovative two-stage cGAN structure that allows diverse, authentic, and flexible subtype-conditioned FOG data generation, and demonstrate its potential for personalized augmentation. To address the unexplored impact of DL-based augmentation on DL FOG detection performance, we evaluate Hi-CF cGAN using a CNN model and compare it against classical augmentations.

## II. Materials and Methods

### A. Dataset and Data Preprocessing

We used the 3D accelerometer signals from both ankles (256 Hz) of a previously collected dataset by Cockx et al. [32] [33]. After excluding one of the participants (Participant 8) due to a misdiagnosis of PD, 15 patients (11 male, 4 female; mostly aged 60–79) and a total of 8.6 hours of data remained. Each participant performed (in OFF medication state) four to six rounds of the same FOG-provoking and real-life-inspired protocol, lasting approximately six minutes each. A round consisted of standing-in-place turns, narrow pathway navigations, straight walking, stopping and starting, turning, and doorway navigation. Labels for FOG (20.11%) and its subtypes (4.14% shuffling, 88.28% trembling, and 7.58% akinesia) were manually annotated by experts.

The dataset was split into 10 round-independent folds using balanced fold assignment algorithm [6]. This algorithm allows each fold to have similar sample counts for non-FOG, FOG, FOG subtypes, and long FOG (>4s) to ensure data consistency and representativeness across folds. An 80%(8-folds)-10%(1-fold)-10%(1-fold) train-validation-test split was conducted. In each iteration of 10-fold cross-validation, the roles of folds rotated sequentially. Raw acceleration signals were then segmented into 2-second windows with 0.5-second overlap, covering at least one gait cycle of PD patients (average stride time 1.12 ± 0.12 seconds [34]). A window was labeled FOG if over 50% of it overlapped a FOG episode or if it fully contained a brief (<1s) episode.

### B. Hierarchical Coarse-to-Fine cGAN (Hi-CF cGAN )

Hi-CF cGAN is made up of a generator (G) and four same-architecture discriminators 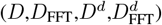, respectively visualized in Fig. 1 and 2.

**Fig. 1:**
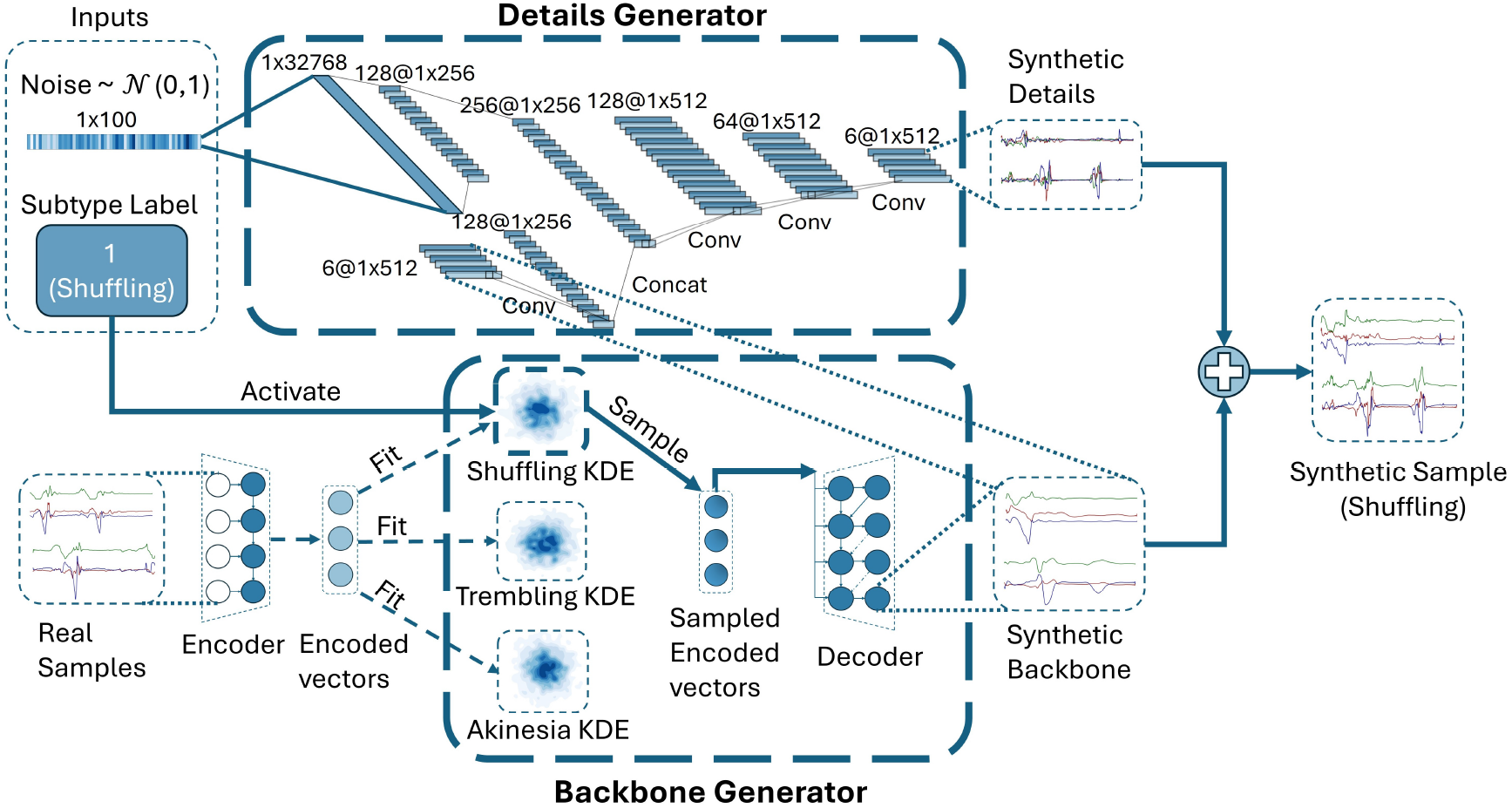
Generator architecture. The generator comprises a backbone generator (*G*^*b*^) and a details generator (*G*^*d*^), which respectively synthesize the coarse and fine-grained patterns with subtype awareness. *G*^*b*^ includes encoder, decoder, and subtype-specific KDEs pre-trained on real signals. A subtype label activates the corresponding KDE to sample an encoded vector, which is then decoded into the synthetic backbone. *G*^*d*^, a parallel CNNs, takes the synthetic backbone and random noise as input to produce synthetic details. Adding synthetic details and backbone yields the final synthetic signal.

**Fig. 2:**
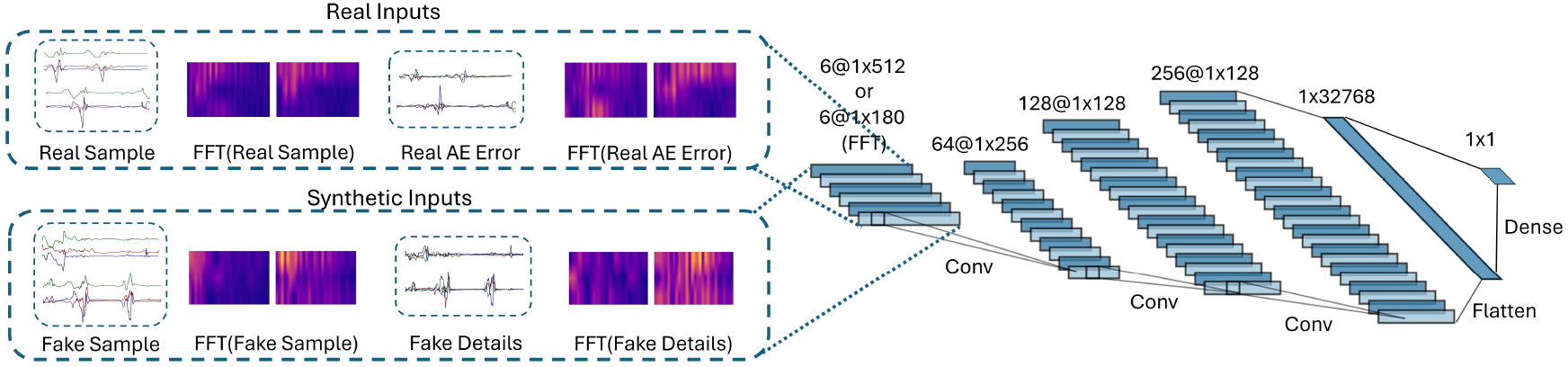
Discriminator architecture. A three-layer CNN that takes real and fake full signals, details, or their frequency representations as input and outputs a value representing real-vs-fake discrimination.

#### 1) Generator Architecture

Our two-stage generator in Fig. 1 produces synthetic samples by combining the outputs of two components: a backbone generator (*G*^*b*^), which generates coarse signal pattern (general trend); and a details generator (*G*^*d*^), which adds fine-grained fluctuations around the general trend. *G*^*b*^ promotes diversity in synthetic signals by introducing variations in general motion trend, enabling both signal generation that captures different motion subtypes with natural intra-subtype variability, and controlled synthesis based on specific motion trends, beyond the classical signal generation driven by noise. With *G*^*b*^ handling the overall motion, *G*^*d*^ can operate in a more focused context, generating sharper and more realistic local artifacts. Although the two generators are functionally distinct, they interact through architecture design and joint training to produce coherent outputs. As auxiliary modules, an Autoencoder (AE) and multiple Kernel Density Estimators (KDEs) are fitted preceding cGAN training.

The AE comprises an encoder and decoder. The encoder processes a raw signal through a 2-layer, 128-unit Long Short-Term Memory (LSTM) to extract temporal information, which is compressed into a lower-dimensional (1×256) encoded vector via a dense layer. The decoder mirrors this structure to reconstruct inputs from encoded vectors. Supporting model training and evaluation, the AE extracts the real backbone, details, and encoded vectors, respectively the reconstruction, reconstruction error, and encoder output of real signals. It also helps construct generated backbone and provides reconstruction of generated sample.

The backbone generator consists of three KDEs, each non-parametrically modeling the distribution of real encoded vectors of one subtype. A subtype label input activates the corresponding KDE, which samples an encoded vector that the decoder reconstructs into a generated backbone.

The details Generator has a parallel architecture. One branch expands Gaussian noise input via a dense layer. The other branch processes the synthetic backbone from *G*^*b*^ as a conditioned input through convolution. The outputs from both branches are concatenated and pass through three 1D convolutional layers, which progressively reduces dimensionality, producing the final generated details matching the sample size.

#### 2) Discriminators Architecture

The discriminators process different inputs while sharing the same architecture (Fig. 2), comprising three 1D convolutional layers, a dense layer, and an output neuron producing maximally distinct values for real and generated inputs. Discriminator D distinguishes the real and fake full signals in time domain (TD), while discriminator *D*_FFT_ differentiates their frequency domain (FD) representations, i.e., magnitude spectrums of Fast Fourier Transform (FFT). Similarly, discriminator *D*^*d*^ and 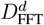 respectively separate the real and generated details and their FFTs. Capturing both TD and FD characteristics is essential since FOG exhibits disrupted temporal [35] and frequency [2] dynamics. Previous studies also found discriminator intaking solely raw data insufficient for generating realistic frequency patterns [29].

#### 3) Objective Functions

The generator loss,*L*_G_, defined in, is a weighted sum of adversarial components 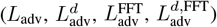for ensuring realism, diversity terms 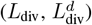 to encourage variations of generated samples, and a reconstruction loss component (*L*_rec_) to achieve a structurally and semantically coherent integration of the generated backbone and details. *α*_1_, *α*_2_, *α*_3_, *α*_4_ were empirically set to 0.5, 0.5, 1, and 0.25, respectively.

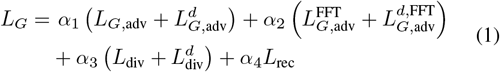

Each adversarial component is computed as the negative of batch-wise average of discriminator outputs on generated samples, following the Wasserstein GAN framework [36]. A higher discriminator output indicates a larger deviation from the real samples. 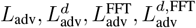 are calculated from the outputs of *D, D*_FFT_, *D*^*d*^, 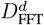 respectively, each evaluated on full generated signal, FFT of full generated signal, generated details, and FFT of generated details. The diversity term is defined as the negative of the average of pairwise Euclidean distances between generated samples within a batch, calculated separately for the full generated signal as *L*_div_ and the generated details as 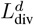. The reconstruction loss is defined in (2) as the average of weighted basic reconstruction loss, *l*, clamped with *M* . *τ, ω*, and *M* are set to 0.1, 5, and 3000 empirically. *l*, shown in (3), is based on Δ_*i,t,d*_, the absolute difference between generated backbone and reconstruction of full generated signal, with stronger penalties for larger difference. The weight factor of *l* is proportional to the gradients of reconstruction 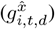 and generated backbone 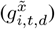, boosting *l* in regions with high temporal dynamics to better capture temporal transitions.

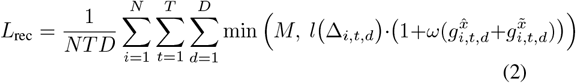

where *N* is the batch size, *T* is the total number of time steps, and *D* is the number of accelerometer channels.

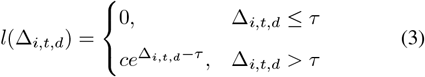

The loss of each discriminator is the sum of an adversarial term and a gradient penalty. The adversarial term is computed as the difference between the batch-wise average of discriminator outputs evaluated on generated and real inputs. A larger difference indicates stronger discriminative ability. The gradient penalty follows the implementation proposed by Gulrajani et al. [37] (*λ* = 10, *β ∼ U* (0, 1)), using interpolated samples between real and generated inputs to promote training stability. For *D,D*_FFT_,*D*^*d*^, 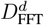, the corresponding real inputs are the real full signal, its FFT, the real details (i.e., reconstruction error of real signal), and the FFT of real details; the generated inputs follow the same structure.

### C. Model Training

Only FOG samples from the train set were used in our cGAN training. The initial learning rate was 0.0001. RMSprop optimizer was used for updating the parameters of the generator and discriminators. The generator and discriminators were trained alternately, following the standard GAN training procedures [38]. At the beginning of each epoch, FOG subtypes were balanced via oversampling to ensure that each subtype was equally learnt. After that, for each real batch, the generator created a same-size synthetic batch conditioned on the subtype labels of the real patch, so that both batches share the same subtype composition. The discriminators were then updated based on their respective losses. Subsequently, the generator was updated using its loss function based on the outputs of the updated discriminators. Through this adversarial process, the generator learnt to produce increasingly authentic signals by learning to deceive the discriminators. Simultaneously, the discriminators refined its ability to distinguish synthetic from real samples, encouraging continuous further improvement of the generator. The training stop criterion was based on the stabilization of *L*_*G*_. If the percentage change of the epoch-wise average of *L*_*G*_ was smaller than a threshold *ε* (*ε* = 0.1) for a maximum of *s* consecutive epochs (*s* = 5), the training was terminated.

### D. Model Evaluations

#### 1) Qualitative Assessments

By direct visualization, real full signals, their FFTs, backbones, and details were compared to those of generated signals to examine the authenticity of generated data and verify model design. Additionally, we visualized Uniform Manifold Approximation and Projection (UMAP), a dimension-reduction algorithm that preserves both local and global data relationships, to evaluate the authenticity and diversity of generated samples and their properties by comparing their distributions to the real ones from the train set. UMAPs were plotted with real and generated signals, their FFTs, and extracted hand-engineered features, separately for each subtype. The hand-engineered features included 13 time domain features (mean, standard deviation, median, trimmed mean, root mean square, median absolute deviation, 25th percentile, 75th percentile, interquartile range, skewness, kurtosis, normalized signal magnitude area, and mean crossing rate) and 3 frequency domain features (Spectral entropy, freezing index, and peak frequency) typically used in FOG detection studies [6] [39]. These features were computed separately for the six accelerometer channels, yielding 78 time domain and 18 frequency domain features. Among these, the vertical-acceleration-based FI [15], the most interpretable and prominent feature for FOG characterization, was specifically plotted.

#### 2) Quantitative Assessments

To quantitatively evaluate the generated samples, we used the unbiased Maximum Mean Discrepancy (MMD), a kernel-based non-parametric measurement of discrepancy between two distributions that supports unequal sample sizes. A smaller MMD indicates greater similarity between distributions. We used Gaussian Radial Basis Function kernel with width set to the median pairwise sample distance. Real-Real MMD was computed between seen (train set) and unseen real samples (test and validation sets), while Generated-Real MMD was computed between generated samples (matched in size to the train set) and unseen real samples. Comparing these two MMDs provided a quantitative assessment of the diversity and authenticity of the generated samples relative to realistic variability. MMDs were computed independently for each subtype on raw signals, their FFTs, and hand-engineered features.

### E. Proposed Augmentation Application in FOG Detection

To assess the practical impact of Hi-CF cGAN augmentation on the performance of a DL FOG detection model, binary classification CNNs, sharing the discriminator architecture plus a sigmoid output activation, were trained with the original dataset and different augmented train set. We used Adam optimizer and binary cross-entropy loss. During training, at the beginning of each epoch, we under-sampled non-FOG of the train set to match the FOG sample size. We applied early stopping with a patience of 10 epochs, based on the losses on the original validation set, to prevent under- or overfitting. Performance of a CNN was evaluated via its detection rates of FOG and non-FOG on the original test set, with FOG detection further broken down by subtype accuracy.

#### 1) General Augmentation

To evaluate the effectiveness of HiCF-cGAN without altering subtype distribution, CNNs were trained on train set modified via general augmentation, i.e., applying the same HiCF-cGAN augmentation ratio (the ratio of real to augmented data) to all subtypes. To examine the impact of augmentation ratios, ratios of 1:0 (no-augmentation), 1:0.5, 1:1, and 1:2 were tested.

#### 2) Comparison to Classical Augmentations

To compare Hi-CF cGAN with classical augmentations, we followed the implementations by Um et al. [28] for rotation, jitter, scaling, permutation, magnitude warp, time warp, and a combination of rotation, permutation, and time warp, which is the most effective mixture augmentation they reported. Using the optimal augmentation ratio from Sec. II-E.1 for all classical augmentations, separate CNNs were trained on train sets modified by each classical augmentation.

#### 3) Personalized Augmentation

Another application of interest was personalized augmentation, where subtype augmentation ratios were adjusted based on the patient’s specific subtype composition. To explore this, we selected two participants with different predominant minor subtypes: Sub-01 (shuffling-dominated) and Sub-15 (akinesia-dominated). For each subject, his or her data served as the test set, while the data of the remaining subjects was used for training and validation, on which AE, KDEs, and Hi-CF-cGAN were trained. Then, a CNN classifier were trained with no augmentation, general augmentation, and a customized augmentation ratio, and their performances were compared.

## III. Results

### A. Qualitative Assessment

Fig. 3, 4, and 5 respectively display the representative generated signals of shuffling, trembling, and akensia and their closest real counterparts (based on backbone similarity using Euclidean distance) in the train set. The signal components (backbones, details) and FFTs are also displayed.

**Fig. 3:**
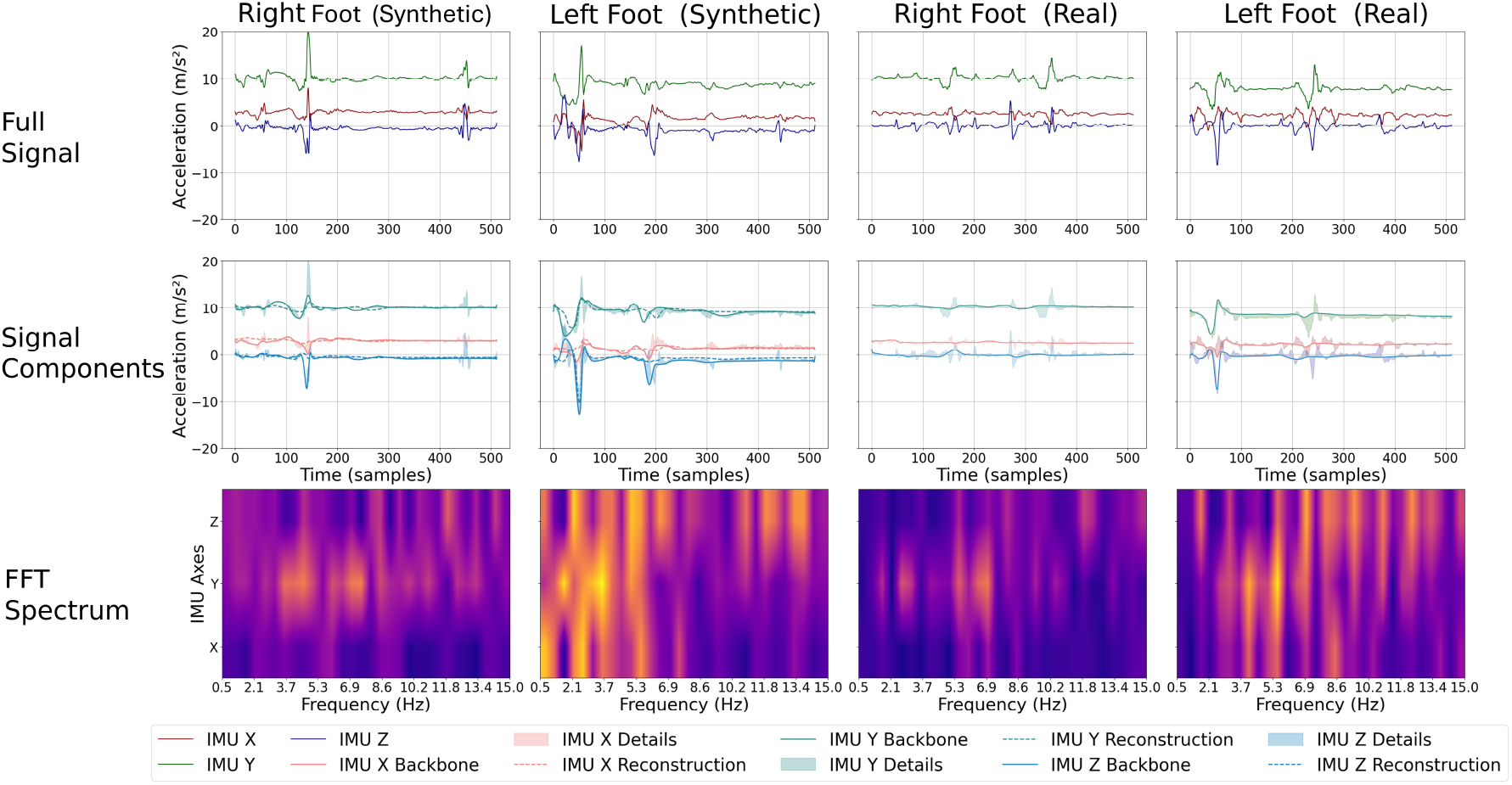
Representative generated and real shuffling signals, demonstrating similar TD and FD characteristics.

**Fig. 4:**
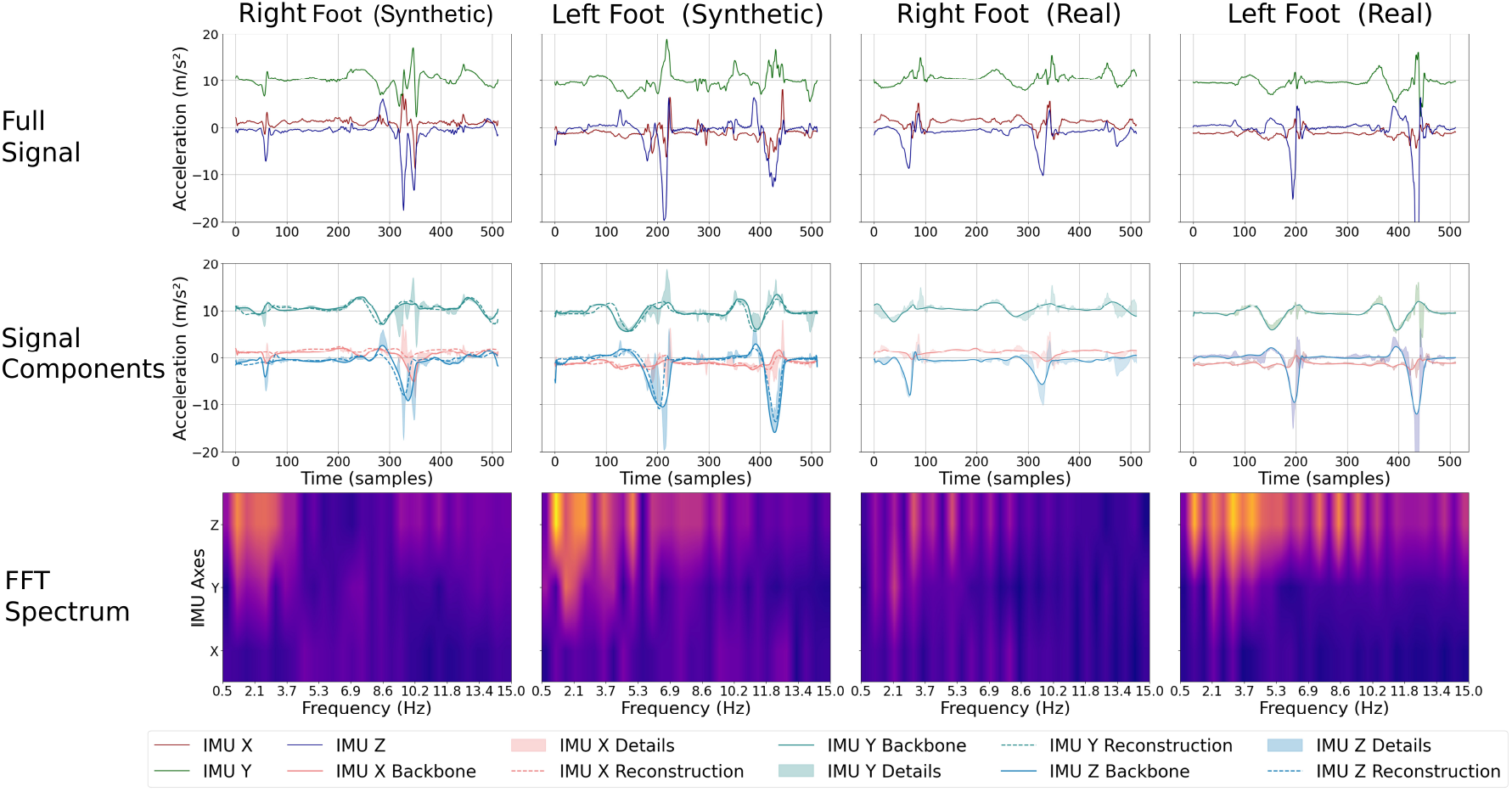
Representative generated and real trembling signals, demonstrating similar TD and FD characteristics.

**Fig. 5:**
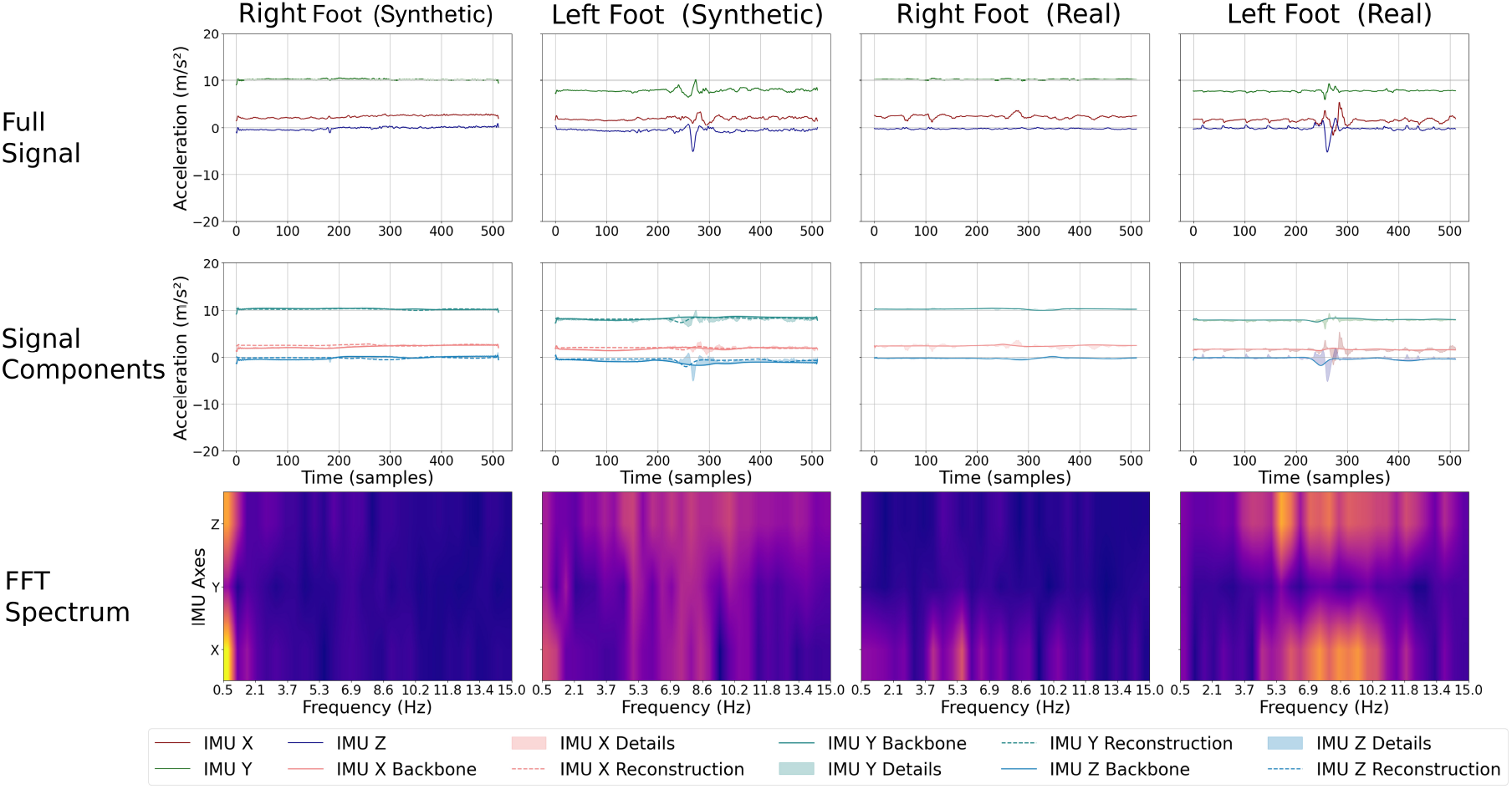
Representative generated and real akenisia signals demonstrating similar TD and FD characteristics.

According to the first rows of Fig. 3-5, the full generated signals appear realistic to human perception based on their plausible channel ranges and realistic noise characteristics with respect to the real signals for all three subtypes. For the motion-heavy subtypes, shuffling and trembling, the intuitive authenticity is also evidenced via their observable gait-like patterns or alternating left- and right-foot movements. Upon closer inspection in the second rows of Fig. 3-5, the generated backbones present a natural global motion trend via their gait-like fluctuations within ranges similar to the real ones. The generated details resemble genuine noise patterns by amplifying near backbone peaks and diminishing elsewhere, coherently combing with the backbones. Additionally, a close match of the reconstructions of the generated signals and their generated backbone indicates the desired coherent integration of details and backbone as *L*_rec_ enforces. The generated and real FFT spectrums in the third rows of Fig. 3-5 also exhibit similar patterns for all subtypes, despite slightly higher concentration at the lowest bar for akinesia.

Fig. 6 visualizes the UMAPs on the real and generated signals, their FFTs, hand-engineered features, and distributions of FI for all subtypes. The real signals are from the train set of a randomly selected fold. For visual clarity, 300 generated signals are plotted on each UMAP, and the highest 5% of FI values are omitted in each FI plot.

**Fig. 6:**
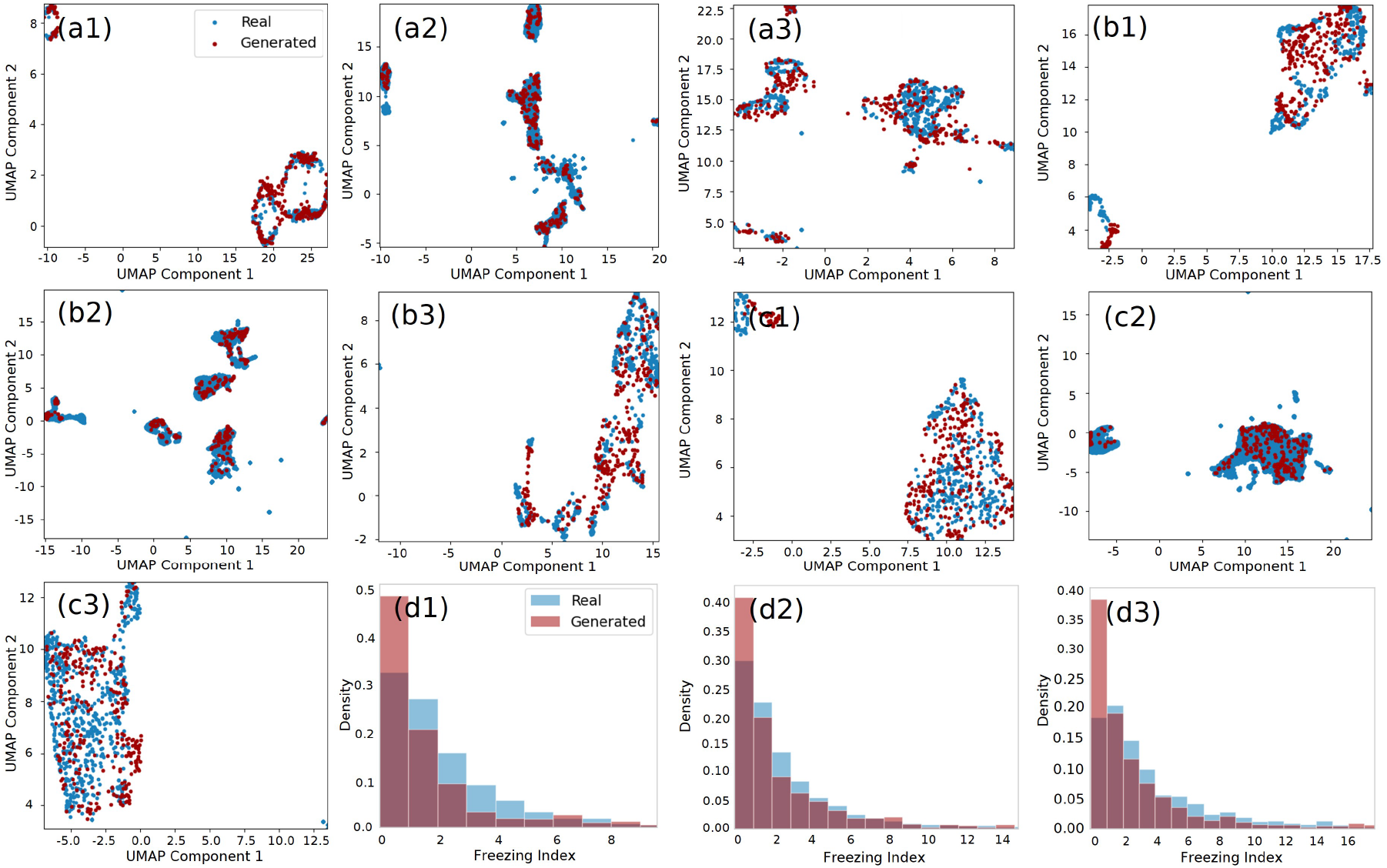
UMAPs on (a) raw signals, (b) FFT of signals, (c) hand-engineered features, and (d) FI distributions for shuffling (1), trembling (2), and akinesia (3). Real and synthetic data show closely aligned distributions across all UMAPs and FI plots.

The distributions of real and synthetic data closely align in all UMAPs in Fig. 6. The generated data spans broadly across the real distribution without local clustering. They also tend to fill up gaps within real data and expand slightly beyond the real distribution boundary. Across subtypes, trembling exhibits the most complete real data coverage. Shuffling shows a minor mismatch with the isolated small cluster in FFT (b1) and hand-engineered features UAMP (c1). Upon closer inspection, that small cluster corresponds to a subject whose left foot sensor was intermittently off. The generator was able to generate in similar channel ranges but deviated more from the corrupted signal patterns due to limited reference samples. Since this discrepancy corresponds to occasional anomalies, it is outside the scope of intended improvements. Akinesia exhibits slightly sparser overlap at the central region in raw window (a3) and hand-engineered features UMAP (c3). Further examination does not reveal any systematic difference between the generated and real data in those regions.

All FI distributions of real and generated data in (d) display a similar general pattern with greater concentration at the lower FI range, despite the generated data exhibiting higher density at the lowest density bar (FI < 1). The FI ranges in generated data (0.00723 to 9.78, 0.0306 to 15.1, 0.00761 to 17.4) matches those in real data (0.00859 to 9.54, 0.00855 to 15.1, 0.0176 to 18.4) across shuffling, trembling, and akinesia.

### B. Quantitative Evaluation

The 10-fold Real-Real and Generated-Real MMDs of raw signals, their FFTs, and hand-engineered features for all subtypes are visualized in Fig. 7. A two-sided Wilcoxon signed-rank test with Bonferroni correction factor of 3 is conducted between each Real-Real and Generated-Real MMD pair.

**Fig. 7:**
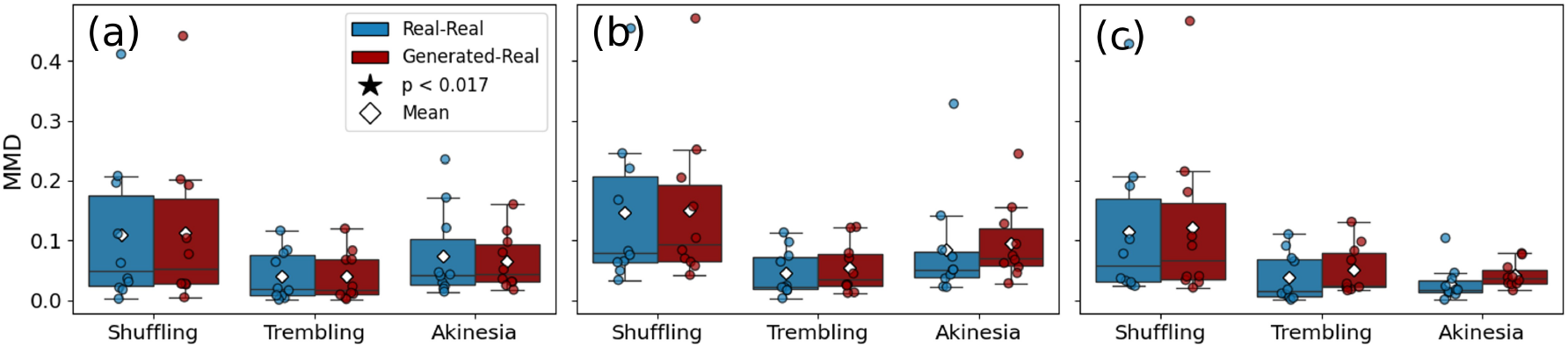
Real-Real and Generated-Real MMDs across subtypes for (a) raw signals, (b) their FFTs, and (c) hand-engineered features. No statistically significant differences were found between any MMD pairs.

According to Fig. 7, no statistically significant difference is found between any Real-Real MMD and its corresponding Generated-Real MMD. Within each pair, although largely comparable, the average Generated-Real MMD tends to be slightly higher than the average Real-Real MMD, which is most noticeable for hand-engineered features and least evident for the raw signals. Across subtypes, the 10-fold coefficient of variation (CV) of MMD is the highest for shuffling (0.876 to 1.195) and similar for trembling (0.781 to 1.068) and akinesia (0.504 to 1.082), while typically smaller for akinesia despite its higher upper bound.

### C. General Augmentation Effectiveness

Fig. 8 shows the 10-fold detection rates of non-FOG, FOG, and FOG for each subtype (via post-analysis) of CNN classifiers trained with general augmentation with different augmentation ratios. For each metric, a two-sided Wilcoxon signed-rank test with a Bonferroni correction factor of 3 is conducted between the baseline and each augmentation ratio.

**Fig. 8:**
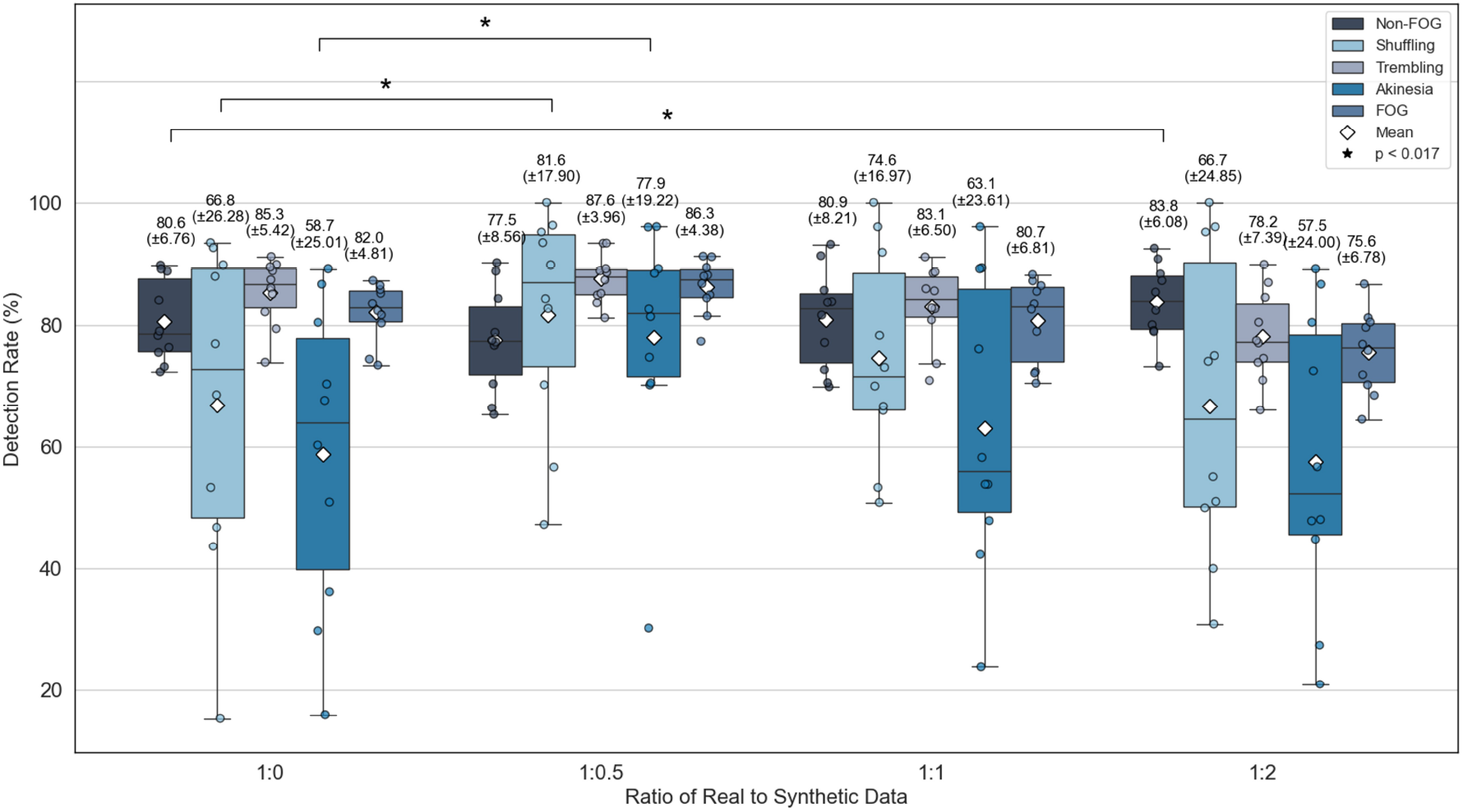
10-fold detection rates of CNN classifiers with general augmentation via Hi-CF cGAN at different augmentation ratios. Mean (standard deviation) values are shown above each group. Minor subtypes (shuffling and akinesia) show significant enhancements at 1:0.5 ratio.

Fig. 8 presents that the detection rate enhancement is initially on FOG-related metrics, especially the minor subtypes, and then gradually shifts onto non-FOG as the augmentation ratio increases. With 1:0.5 augmentation, the detection rate is significantly improved in shuffling (p = 0.00781) and akinesia (p = 0.00391) compared to the baseline (1:0) augmentation. Moreover, the 10-fold detection rate variations of FOG and all subtypes are reduced; and the mean detection rates of trembling and FOG are also boosted. With 1:1 augmentation ratio, although not statistically distinguishable, the mean detection rates of shuffling and akinesia also improve. With 1:2 augmentation ratio, a statistically significant improvement is found with non-FOG (p = 0.00977). However, with our primary focus on FOG detection, we select 1:0.5 as the optimal augmentation ratio for the following evaluations.

### D. Hi-CF-cGAN compared to Classical Augmentations

Fig. 9 displays the 10-fold average detection rate of FOG, non-FOG, and each subtype of CNNs trained with no augmentation (baseline), Hi-CF-cGAN-based augmentation, and classical augmentations.

**Fig. 9:**
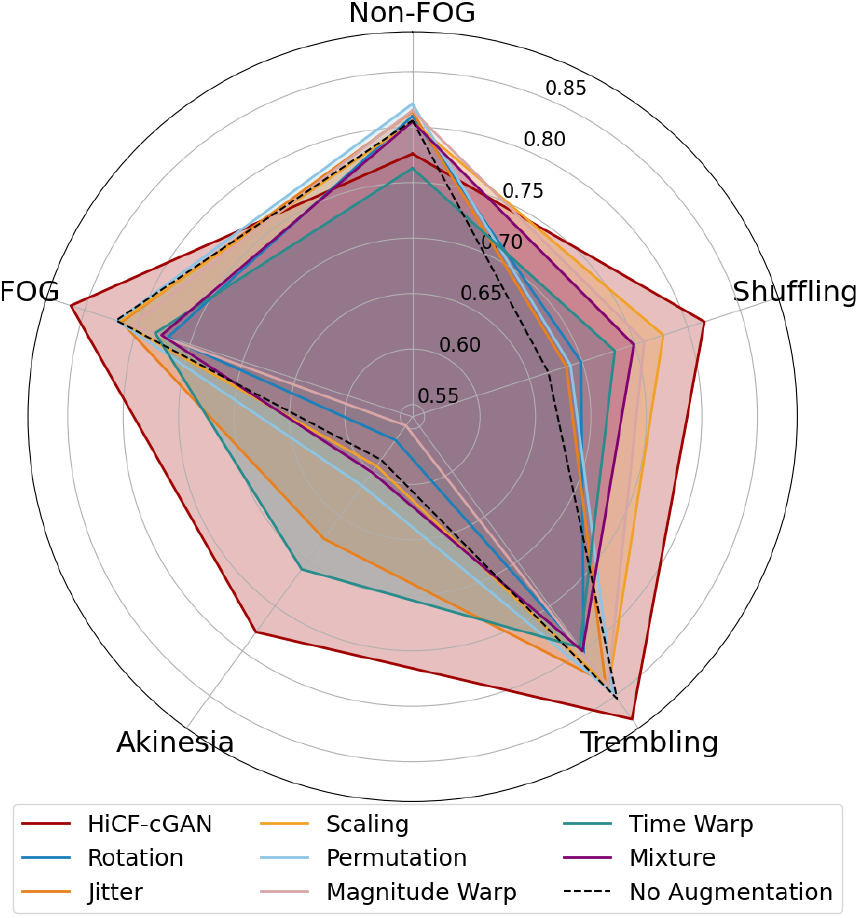
10-fold average detection rates of CNN classifiers with general augmentation via Hi-CF cGAN and classical augmentations. Hi-CF cGAN achieves the greatest and most comprehensive improvements over baseline for FOG detection overall and across subtypes.

Presenting a markedly distinct contour in Fig. 9, Hi-CF-cGAN-based augmentation leads to the highest improvements from the baseline for FOG detection, both in general and for all FOG subtypes, most effectively resolving the original imbalanced subtype performances, with a minor degradation in non-FOG detection rate (-3.09%), which is smaller than the gain in FOG detection (+4.24%). Additionally, Hi-CF cGAN achieves the most comprehensive FOG-related improvements without the trade-offs between metrics observed with classical methods. While classical augmentations yield improvements in either shuffling subtype (using rotation, magnitude warp) or across both minor subtypes (using jitter, permutation, scaling, time warp, and mixture method), these gains are consistently accompanied by performance degradation in FOG and trembling and occasionally in akinesia or non-FOG metrics and are smaller than those by Hi-CF cGAN. Leveraging two-sided Wilcoxon signed-rank tests, the enhancements by Hi-CF cGAN are significantly higher than those by classical augmentations in shuffling (p = 0.00769-0.0382), akinesia (p = 0.00769-0.0117), and FOG (p=0.00391-0.0371). The only exceptions are classical augmentation by scaling for shuffling, and augmentation by time warp for akinesia. In trembling, Hi-CF cGAN exhibits significantly higher gain compared to time warp (p = 0.0273) and mixture method (p = 0.00586). The remaining advantages in trembling for Hi-CF cGAN compared to classical augmentations are also remarkable.

### E. Personalized Augmentation Effectiveness

The FOG episodes captured from Sub-01 consist of 82% of trembling, 16% of shuffling, and a negligible amount of akinesia (2%). Since trembling is the dominant subtype, consistent with the general dataset trend, the customized augmentation ratio for this subject is set 1:0.5 for shuffling only. In contrast, Sub-15 exhibited a balanced distribution of 49% trembling and 51% akinesia. Given the comparable prevalence of these two subtypes, both were augmented at a ratio of 1:0.5. The performance of CNNs trained with no augmentation (baseline), general augmentation, and personalized augmentation are shown in Fig. 10.

**Fig. 10:**
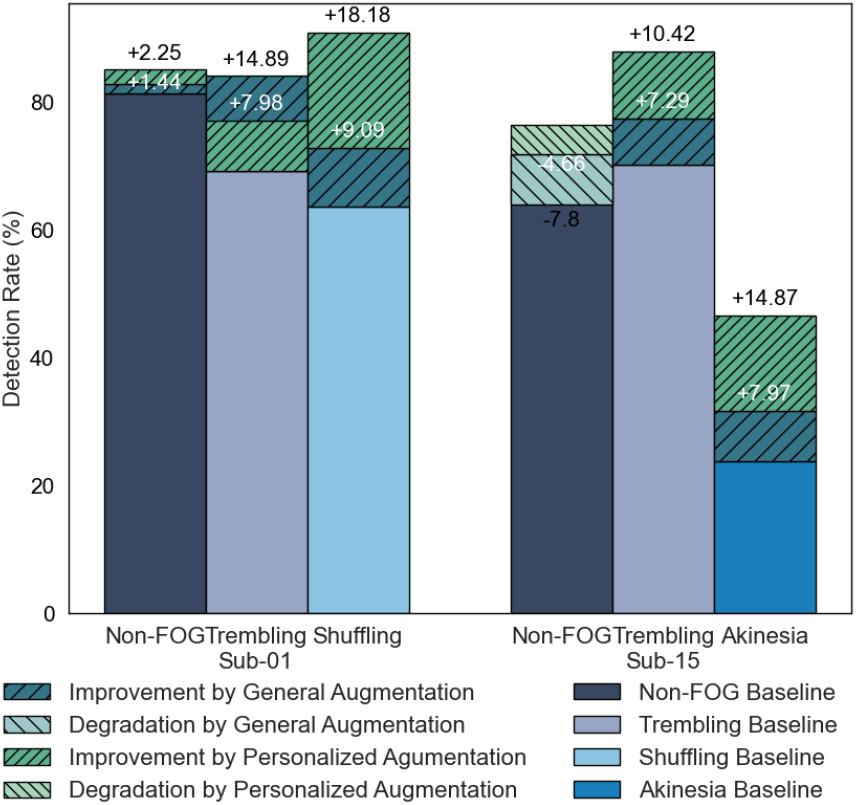
Detection rates of CNN classifier with different Hi-CF cGAN augmentation settings. Personalized augmentations improve baseline performance more than general ones.

According to Fig. 10, for Sub-01, both the general and customized augmentations improve detection rates of non-FOG (+1.44%, +2.25%), trembling (+14.89%, +7.98%) and shuffling (+9.09%, +18.18%) from the baseline. The personalized augmentation leads to higher enhancements for non-FOG and especially shuffling, the original minor subtype, than the general augmentation. For Sub-15, both the general and personalized augmentation improve both trembling (+7.29%, +10.42%) and akinesia detection rates (+7.97%, +14.87%). Although a non-FOG degradation (-7.8%, -4.66%) is associated with the enhancements, it is smaller than the overall FOG gain based on average subtype gain. The customized augmentation also shows higher improvements on both subtypes and less cost in the non-FOG compared to the general augmentation.

## IV. Discussion

Authenticity and diversity are the principal evaluation criteria for generated data. For all subtypes, the authenticity of the FOG signals generated by Hi-CF cGAN, with respect to real signals, is supported by (1) direct visual comparisons; UMAP-revealed and MMD-evaluated distributional consistency in time-domain, frequence-domain, and hand-engineered features (not directly optimized); and (3) alignment of distributions and ranges of FI. The diversity of the generated signals is qualitatively evidenced through its broad and non-clustered UMAP distributions, indicating a sufficient coverage of the variance of real distributions; and through its gap filling and broader extension compared to the real distributions, expanding the original dataset coverage and thus enhancing the generalizability. Quantitatively, the averages of Generated-Real MMDs, which reflect the differences between the distributions of synthetic and real data, tend to be higher than those of the corresponding Real-Real MMDs, which represent the intrinsic variability within the real data. Given the lack of statistically significant differences across all MMD pairs, this elevated discrepancy remains within a reasonable range and is interpreted as an indication of improved generative diversity rather than unrealistic deviation from the real data distribution.

Despite the general realism and richness of the generated samples, their quality varies slightly across metrics. Based on the differences between average Generated-Real and Real-Real MMDs, the authenticity was the highest with raw signals, exhibiting the lowest difference, followed by its frequency-domain representations, and the lowest with hand-engineered features. In contrast, diversity is the highest in hand-engineered features, then frequency-domain representation, and lowest in raw signals. This may be explained by the intrinsically higher sensitivity of frequency-domain information to subtle signal variations compared to the raw signal in time-domain, which could average out the minor differences across its large number of time points. The higher variability in hand-engineered features is expected since the discriminators did not directly target hand-engineered features via their inputs or loss functions. As an example, the distribution of FI of the generated data demonstrates higher density in the lowest frequency range, in line with previous study [31]. This slight bias in characteristic feature is due to the exaggeration of the real frequency-domain trend targeted by*D*_FFT_ and 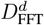, which favors lower frequency components. It is not corrected, since FI is not explicitly targeted by any discriminator, unlike the general frequency-domain representation.

Moreover, the generative quality evaluation varies slightly across subtypes. With its most thorough UMAP real coverage, generated trembling signals show better authenticity performance than shuffling and akinesia. For shuffling, a partial mismatch was observed in an outlying UMAP cluster. However, as noted in the Sec. III-A, this stems from a rare signal corruption caused by a dysfunctional left foot sensor and is therefore outside the model improvement scope. For akinesia, the sparse overlap between real and generated samples in the central areas of their UMAP projections suggests that real samples there are closer to each other than to the nearest generated samples. This does not imply a failure to capture characteristics unique to the real samples in the above-mentioned UMAP central regions, since no systematic differences were found between real and synthetic samples there. Instead, this suggests that real akinesia samples are intrinsically more tightly clustered and thus more sensitive to modifications. This also explains the higher low-frequency magnitude bar in the direct FFT visualization of generated akinesia signals, which arises from minor high-frequency fluctuation amplified by the overall quite data. This is consistent with akinesia’s inherent low variance based on its definition and exhibited smallest MMD variation. Consequently, generating authentic akinesia samples is harder than other subtypes under the same evaluation criteria. We thus believe the lower authenticity performances of akinesia reflect partially its phenotype nature and may not necessarily imply a lower realism.

Leveraging the high-quality generated data, subtype-stratified general augmentation effectively improves the CNN performance in detecting FOG, with different augmentation ratios benefiting different detection rates. At the lowest augmented ratio (1:0.5), subtype detection imbalance is reduced, and FOG detection rate is enhanced, with no significant cost at non-FOG detection. This enhancement stems from increased realistic subtype-specific reference samples, which improves discrimination from non-FOG based on subtype-specific characteristics. Additionally, increasing FOG data overall reinforces sharable detection strategies across subtypes. Together, these boost the detection rates of FOG and all subtypes, most significantly in minor subtypes, as they originally lack sufficient reference samples to differentiate from non-FOG. The 10-fold variations in all these FOG-related metrics also decrease, reflecting improved robustness and generalizability towards diverse unseen data of the detection model enabled by added diversity of augmented data. As the augmentation ratio increases, the improvement shifts towards better non-FOG detection. This is because, without data augmentation, the model was slightly underfitted to non-FOG data. This underfitting was a side effect of subsampling to balance the number of FOG and non-FOG samples during training, which forced the model to train on a small, variable subset of non-FOG samples with minimal overlap across epochs. With more FOG data available through synthetic data augmentation and improved class balance, the model could train on larger and more consistent non-FOG samples during training, leading to better generalization and improved non-FOG detection. It is important to realize that even a small improvement in non-FOG detection can already result in a substantial decrease in false alarm rates, as in real-life situations the non-FOG condition is highly more common than the FOG condition. Depending on the application, one might yield more importance to a better FOG detection rate (i.e., sensitivity) versus a better non-FOG detection rate (i.e., specificity).

At the optimal stratified augmentation ratio, Hi-CF cGAN outperforms all tested classical augmentations by simultaneously improving detection rates for FOG and all subtypes and achieving the greatest enhancement extent on each. This can be ascribed to the higher diversity, authenticity, and adaptability of FOG generated data by Hi-CF cGAN. Classical augmentations have lower generative diversity since they only modify original signals to a limited extent. Differently, the diverse backbones of Hi-CF cGAN enforce various motion trends that more likely generalize to unseen samples. Moreover, classical augmentations, based on inflexible hand-crafted rules, offer less assured authenticity. In contrast, guided by adversary training, Hi-CF cGAN demonstrate realistic general trend and details as previously discussed. Additionally, classical augmentations are not customized for FOG signal or subtypes, whereas Hi-CF cGAN implements data-driven and subtype-conditioned generation that adapt to both FOG signal and different subtypes, verified by its comprehensive improvements on FOG and all its subtypes. Without such customizations, only some classical augmentations show notable improvements on some subtype(s) when aligning more closely with some subtype properties. For instance, jittering and time warp improve akinesia detection since they well simulate the minor realistic noise variations across akinesia samples with intrinsically limited fluctuation. Scaling, magnitude warp, time warp, and mixture enhance shuffling detection since they diversify magnitude or temporal gait characteristics, which is suitable for shuffling with clear motion pattern. Trembling, in contrast, was not boosted by any classical augmentations since its underlying highly noisy pattern cannot be effectively diversified by any minor modifications.

Hi-CF cGAN also demonstrates the effectiveness of subtype-variant personalized augmentation, which typically boosts a CNN classifier more effectively than general stratified augmentation by better adapting to individual patient’s needs. We demonstrated the personalization capabilities of Hi-CF cGAN data augmentation on two different patients. In both cases, general and personalized augmentation showed similar improvement trends, enhancing detection across all relevant subtypes with minimal or no degradation in non-FOG detection performance. This similarity is expected, as general augmentation spans the range of the personalized augmentation. However, personalized data augmentation typically yields more pronounced gains by targeting the predominant subtype(s) for each individual. For Sub-01, although only shuffling was augmented, the trembling detection rate also improved, likely due to the frequent co-occurrence and overlapping traits of these subtypes observed in data labeling [33]. Based on the available data, i.e., linear acceleration measured from a sensor affixed above the ankle, our generated data is limited to such accelerations. Future studies may explore other useful kinematic data such as joint angles and angular velocities. Future work may also extend to a more complex setting with more sensors that enable generation of a broader set of body movements. Moreover, expanding augmentation to non-FOG episodes by stratifying between walking, standing, turning, sitting, and other daily-life tasks, may further enhance algorithm performance [40]. Additionally, this study augments FOG episodes to improve detection models. Expanding augmentation to pre-FOG periods may similarly enhance prediction models, enabling earlier and more robust prediction for implementing proactive interventions to prevent FOG or falls.

## V. Conclusion

We proposed Hi-CF cGAN, an innovative two-stage network that flexibly generates subtype-conditioned, authentic, and diverse synthetic ankle acceleration data during FOG. The authenticity and diversity of the generated data, evaluated in the time-domain, frequency domain, and hand-engineered features, were assessed and verified via direct visualization, UMAP projections, and MMD analysis. Evaluation quality was slightly lower in frequency-domain for its sensitivity to modification, and in hand-engineered features due to their lack of direct training focus. Across subtypes, trembling exhibits the highest authenticity while others have some minor while non-significant flaws. Through subtype-stratified augmentation, Hi-CF cGAN successfully improved the detection rates of FOG and subtypes, particularly for previously underrepresented subtypes, in a baseline CNN. Both the comprehensiveness and magnitude of the improvement outperformed those achieved by other classical augmentations, demonstrating the advantageous authenticity, diversity, and adaptability. Additionally, subtype-variant personalized augmentation led to even greater gains by focusing on each individual’s predominant subtype(s), highlighting another promising application. Future work may extend this approach to other kinematic signals, more complex sensor configurations, and pre-FOG generation for prediction applications.

